# Organisation of feed-forward loop motifs reveals architectural principles in natural and engineered networks

**DOI:** 10.1101/188821

**Authors:** Thomas E. Gorochowski, Claire S. Grierson, Mario di Bernardo

**Author notes:** These authors contributed equally to this work (Claire S. Grierson, Mario di Bernardo).

## Abstract

Network motifs are significantly expressed sub-graphs that have been proposed as building blocks for natural and engineered networks. Detailed functional analysis has been performed for many types of motif in isolation, but less is known about how motifs work together to perform complex tasks. To address this issue we measure the aggregation of network motifs via methods that extract precisely how these structures are connected. Applying this approach to a broad spectrum of networked systems and focusing on the widespread feed-forward loop motif, we uncover striking differences in motif organisation. The types of connection are often highly constrained, differ between domains, and clearly capture architectural principles. We show how this information can be used to effectively predict functionally important nodes in the metabolic network of *Escherichia coli*. Our findings have implications for understanding how networked systems are constructed from motif parts and elucidates constraints that guide their evolution.

---

Networks are commonly used to represent the complex interactions between components found in natural and engineered systems. Making sense of these structures has so far relied on the analysis of global topological features such as degree distributions or clustering coefficients [1], and the classification of significant localised structures called network motifs [2, 3].

Major progress has been made in the literature towards understanding how some motifs contribute to network structure and function. This has involved proving that motifs exist, are not there by accident, and that they make significant functional contributions to networks [2, 3, 4, 5, 6, 7]. Important families of motifs have been discovered that are shared by diverse networks carrying out similar functions [2, 5] and attempts have been made to relate motif structure with motif function. This has shown that motifs play an important role in gene regulation [3, 8], accelerated response times [9], dynamic stability [10], and responses to noise [11].

Even with these detailed studies, the functional importance of motifs is often uncertain and contested [12, 13]. In particular, it is not clear to what extent the functions of the motifs depend on the context in which they are found i.e., their specific dynamical parameters or their position and connections within the network [13]. For example, Burda et al. evolved gene regulatory networks in silico for user defined functions [14]. They found simple network functions (steady states) resulted in the emergence of motifs where each had an isolated function. However, as more complex phenotypes were chosen, the individual role of the emergent motifs became less clear. Instead, motifs acted more like parts in a larger machine and the function of each motif could only be understood in context.

Some efforts have been made to study larger motif-based structures in complex networks with Kashtan et al. developing the concept of network motif generalizations [15]. These assume that a motif can act as a template from which larger network structures can be built, specifically through duplication of nodes and associated edges that share a similar role (e.g., inputs or outputs, see [15] for a formal definition). An example of such a motif generalization is the bi-fan where two input nodes are connected to a set of output nodes whose number may potentially vary. In this case a single bi-fan (two inputs and two outputs) would form the template motif, and variants with larger numbers of output nodes would be classed as generalizations. Generalizations of motifs often maintained the dynamical function of the template motif, and specific examples of multi-input and -output feed forward loops (FFLs) could test signal persistance and enable the temporal ordering of events [15]. While generalized motifs offer a way of classifying families of related motif, this approach neglects the many ways that a given motif can connect within a sub-structure that does not involve duplication, or connections between motifs of completely different types (e.g., feed forward loops connected to feedback loops).

We set out to discover large statistically over represented structures and capture additional building blocks of networks whilst taking into account the diversity of connections between parts. Previous studies have neglected these aspects. Our understanding of how motifs distributed through a network co-ordinate and tune their collective function through different types of connection is therefore limited. This is further hindered by current analysis methods being fundamentally unable to capture how localised network motifs are connected to produce functionally important topological features at these intermediate scales. Even though motif aggregation has been observed previously in several biological networks (e.g., gene transcription [16] and protein interactions [17]), and shown to be interwoven with global statistical properties [18], there currently exists no standard way to quantify the spectrum of possible connection types and extract the rules that might underly the aggregation process.

In this work we present new tools that uncover the precise organisation of connections between network motifs in order to detect, categorise and quantify motif clusters. Furthermore, we study how information flows through the nodes in these structures. Focusing on the widespread and highly studied FFL motif, we investigate how structures of FFLs are organised in a range of natural and engineered networks. Our results reveal highly distinctive types of FFL cluster for different types of network. Random networks have very different distributions than the natural and engineered networks that we tested. Even though many types of clustering are possible, often just one or two types dominate, forming over 80% of the FFL clusters. Which cluster types dominate depends on the type of network. We illustrate that characterising network structures at the scale of motif clustering produces highly distinctive and surprisingly simple profiles from which clear conclusions about network structure, function and evolution can be drawn.

## Results

A broad range of biological, engineered and social networks were selected for analysis. These covered the transcriptional regulation of E. coli [4] and S. cerevisiae [2], Gnutella peer-to-peer file sharing [19], Wikipedia voting [20], air traffic control, EU emails [21], Little Rock Lake food web [22], metabolism of E. coli and A. fulgidus [23] and the neural network of C. elegans [24] (see Supplementary Information, Section 2.1 for further details). Although these networks displayed a broad range of global statistics (Supplementary Information, Section 2.1), in every case the FFL motif was found to be significantly over expressed (*p* < 0.0001; Supplementary Information, Section 2.2). This has been recognised previously for many of the networks [2, 5, 4, 7, 23], and suggests that FFLs may play a key functional role across all these systems, acting as a generic building block for many different types of complex system.

## FFLs cluster in real-world networks

Having found FFLs to be a significant feature of the real-world networks, we next investigated their general organisation by studying their propensity to aggregate and become clustered (Fig. 1a). To capture this property we defined a measure of motif clustering (M_c_) calculated as the proportion of shared nodes between all pairs of FFL, normalised by the maximum number of possible shared nodes between all pairs (Fig. 1b; Methods). A motif clustering value of 0 would corresponds to each FFL in our networks being fully isolated (sharing no nodes with any others), while a value of 1 would represent every FFL sharing the same two common nodes with only a single node differentiating each motif (a single fully clustered region). This measure was further generalised to allow for sets containing different types of motif (see Supplementary Information, Secion 1 for an example; Methods).

**Figure 1.**
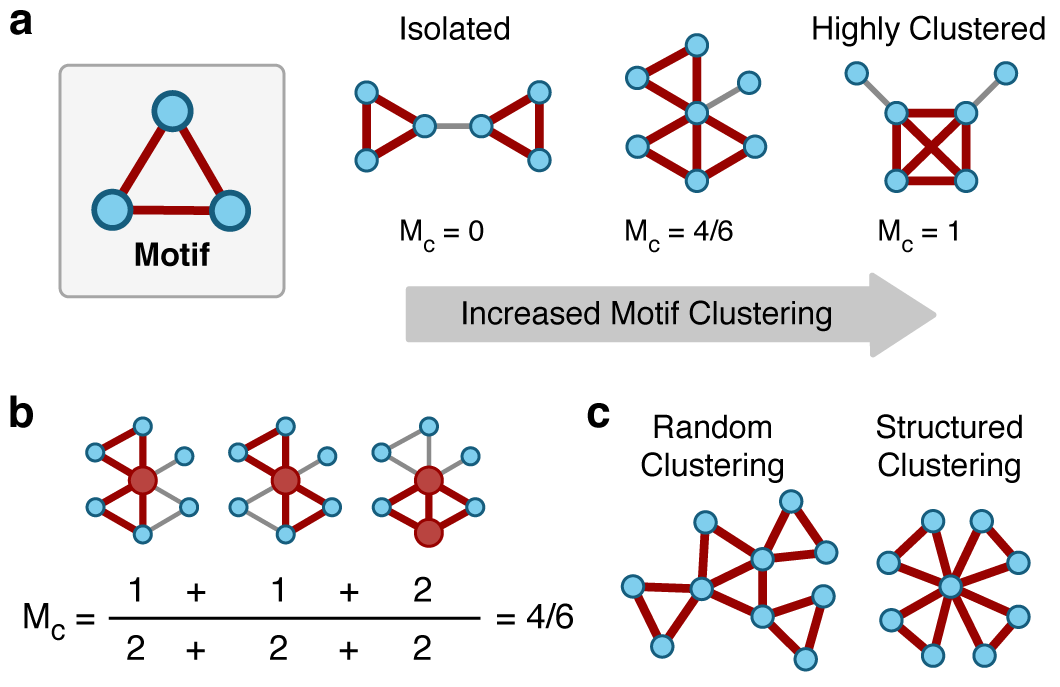
Key features of motif organisation in complex networks. (**a**) The motif clustering coefficient, M_c_, represents the amount of overlap in terms of shared nodes between all pairs of a particular set of motifs. This example shows increasing motif clustering for a set containing a single triangular motif (highlighted in red). (**b**) An example of how the motif clustering coefficient is calculated for the intermediate network from (A). The two motifs and shared nodes have been highlighted in red and for each pair of motifs the maximum possible shared nodes is 2 (found in the denominator). (**c**) Motif clustering can occur in many different ways, being random or structured. Further information regarding specific types of connection between motifs is required if important underlying features are to be understood, see Fig. 2.

Analysis of the networks found FFL clustering to be a significant feature in all the networks, with a positive correlation between overall motif expression and clustering (Supplementary Information, Section 2.2). Increases in motif clustering are inevitable as the density of FFLs increases. However, it is important to note that the significance we report here relates to a null random model that maintains the same number of FFLs (Methods). The motif clustering we see is significantly higher than we would expect given the number of FFLs present.

## Exploring FFL cluster types and their links to network function and evolution

Motif clustering alone told us little about the specific ways in which FFLs had become clustered and whether there existed underlying organisational principles to how they interacted (Fig. 1c). Indeed, the dangers of considering only the global statistical features of a network were highlighted by Li *et al*. [25]. They showed that for a general statistical feature such as the degree distribution, a near identical power law distribution could be generated by completely different types of underlying network structure. Furthermore, specific differences in the way particular nodes were connected in these networks yielded large differences in their performance and reliability to attacks calling into question findings that often state power law and scale-free like networks are inherently susceptible to targeted attacks at hubs. This work clearly illustrated that the differences in local connections really do matter from both a structural and functional perspective.

To ensure that we considered such localised features, we next analysed the specific ways in which motifs were connected by categorising the different possible pairwise combinations of (coherent) FFLs found in our networks, see Fig. 2. These fall into 12 types and we quantified the populations of each FFL combination as a fraction of the total.

**Figure 2.**
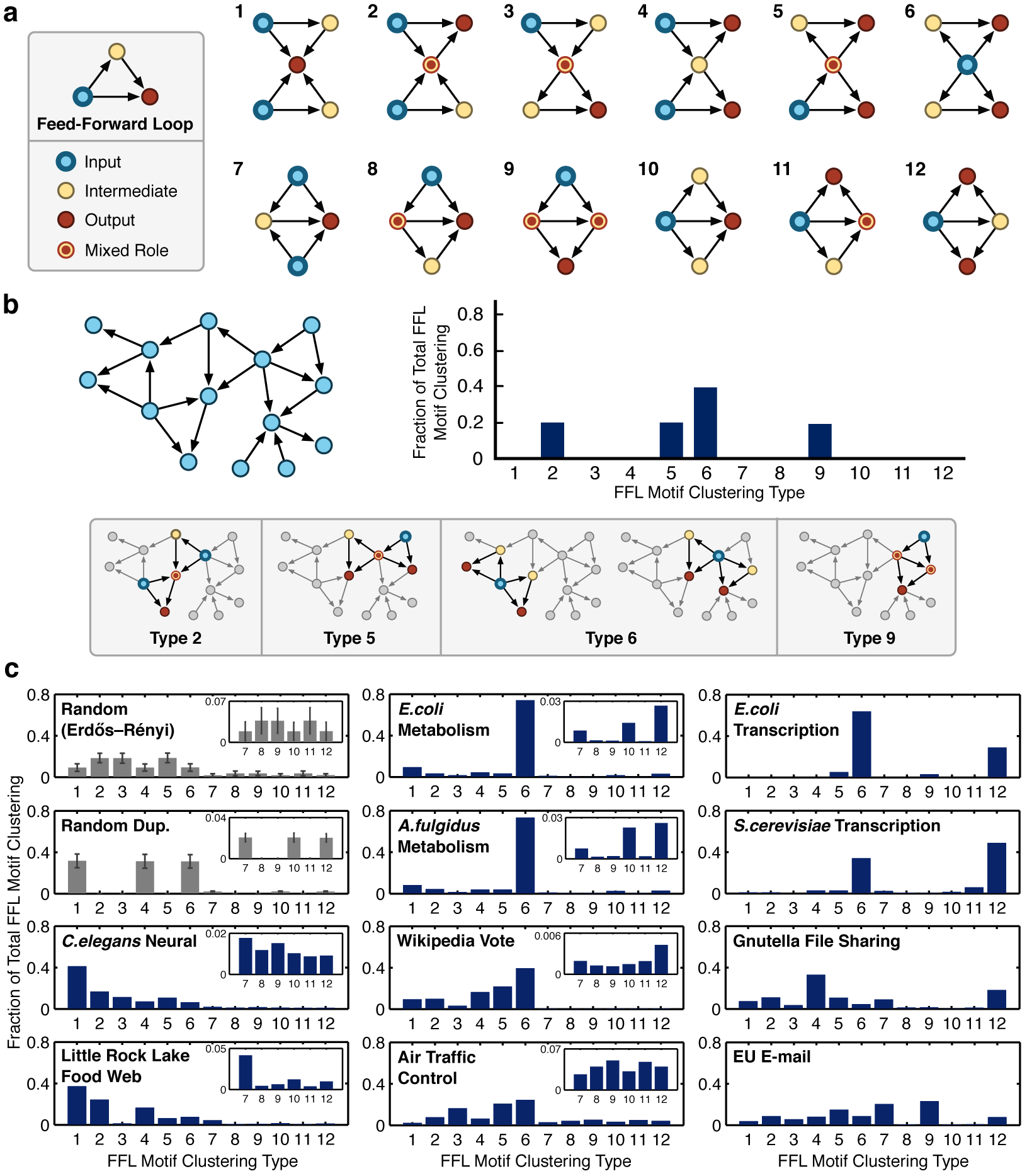
Classifying motif <u>clusteri</u>ng types and their distributions. (**a**) The 12 unique motif clustering types for two feed-forward loops (FFLs). To highlight the flow of information, each node has b<u>een coloured</u> **r**<u>n relation to its role, either: inp</u>ut, intermediate, output, or a mixture of these roles. (**b**) Example network (left) with the associated motif clustering type distribution (right) and the motif pairs and their classifications (below). (**c**) Motif clustering distributions for the natural and engineered networks. Clear signatures are shown in the types of motif clustering found. Inserts have been included for distributions where motif clustering types 7-12 are observed at low overall fractions. The random Erdos-Renyi distribution is generated from a network of 1000 nodes and an edge probability of 0.005 with standard deviations plotted from a sample of 1000 networks. The random duplication distribution is generated from a network of 30 nodes with standard deviations plotted from a sample of 1000 networks.

Looking at the profiles of the natural and engineered networks that we characterised (Fig. 2c), it is apparent that it is not just clustering, *per se*, that matters; different types of FFL cluster are prominent in different networks, and there are similarities between networks with similar functions. This suggests that some types of FFL cluster are better suited to specific functions than others. We investigated this further by focusing on specific types of cluster that are especially over-represented. We hypothesised that these might be cases where one or more specific types of FFL cluster has been subjected to strong positive selection.

## Biological networks

The strongest bias in our data was seen in the two metabolic networks where in both cases over 70% of the FFL clusters are type 6 (Fig. 2c). Type 6 FFL clusters have a single node that is an input to all the other nodes in the cluster (Fig. 2a). In metabolic networks these nodes represent enzymes that produce metabolites that are consumed by multiple enzymes, so the prevalence of type 6 clusters reflects the broad use of the same compound by multiple biosynthetic pathways. It should be noted that the networks we study here include co-factors such at ATP and ADP which are known to bind metabolism together due to their pervasive use and lead to its small-world structure (see Supplementary Information, Fig. S12 for an example of this closely knit architecture with some of the most highly connected node identities included). Furthermore, while the overall concentrations of the combined pools of these co-factors are highly regulated (remaining virtually constant such that individual reactions have little effect), changes in their individual concentrations are known to play important roles in the control of other metabolic functions such as in glycolysis [26]. Therefore, the inclusion of these metabolites was considered important to capture the entire range of potential links within the network.

The functional roles that motifs might play in metabolic networks remain unclear. However, recent comparative studies have found that similarities in the proportions of enzyme classes between species were related to structural features of the motifs present [7]. Comparisons of motif distributions across different organelles within a cell also showed distinct differences which are thought to relate to the specific metabolic functions of each compartment [7]. These differences will directly impact upon the types of motif clustering that are possible and so the highly limited forms we find for FFLs are likely mirrored for other motifs that are found to be prevalent.

Highly constrained motif clustering types were also found for the two transcription factor networks, where two types of FFL cluster, types 6 and 12, make up 80-90% of the total (Fig. 2c). In type 6 clusters genes join with their regulator to co-regulate a single target gene. Type 12 clusters also have a single transcription factor regulating others in the cluster, but in this case the regulator and its target co-regulate multiple target genes. Type 12 clusters enable closer coordination between the expression patterns of groups of genes and have been shown to enable temporal regulation (e.g., ordered activation) of target genes [15].

FFL clusters in both networks display an output centric structure with approximately 60% of nodes acting as outputs. The remaining nodes have an input role, although only a small percentage (12% for E. coli and 16% for S. cerevisiae) act solely as inputs (Fig. 4b). Closer inspection finds that nodes acting solely as inputs are master regulators. For example, in the E. coli network we find the input node CRP (cAMP receptor protein) which has been found to regulate over 180 genes related to catabolism of secondary carbon sources [27] as well as other cellular processes such as biofilm formation, virulence and nitrogen assimilation to name but a few. Another input node FNR, regulates hundreds of genes to control the transition from aerobic to anaerobic growth [28], as well as many other cellular functions.

Another striking feature of these motif clusters is the simple hierarchical structure, known to be present in these types of network [29, 30]. This is most evident in the *E. coli* network where the 7 separated components all share highly similar architectures (Supplementary Information, Fig S3). In this network we tend to find intermediate nodes connected to only a few targets (outputs), as shown in the cluster regulated by CRP. In this case, the large number of separated intermediaries relate to the control of different metabolic processes allowing for a signal from the master regulator (CRP) to be adjusted for a specific processes needs. This separation also accounts for the large numbers of type 6 clustering in the motif clustering type distributions (Fig. 2c). Moreover, the highly specific types of FFL clustering displayed suggests that these structures exhibit important functional benefits, or other possible types of clustering see strong negative selection. The former is supported by dynamical analyses of potential functions such structures might enable [3, 8, 9, 11].

The *S. cerevisiae* motif clusters also exhibit a hierarchical structure, but of a far more integrated form [31] (Supplementary Information, Fig S3) i.e., fewer inputs and intermediaries control many outputs. Lee *et al*. also reported this feature finding that the number of promoter regions bound by a regulator ranged from 0 to 181 with an average 38 interactions per regulator, and many interactions in the same functional category [32]. This is clearly illustrated for the large motif cluster controlled by GLN3 and DAL80 that contains 17 output genes, all regulated by these same two inputs (Supplementary Information, yFig S3). This more integrated architecture leads to an increase in the overall motif clustering shown by the associated z-score (Supplementary Information, Table S2) and helps explain the elevated proportion of motif clustering type 12 where a single input and intermediate are connected to large numbers of outputs (Fig. 2c).

Unlike the *E. coli* networks, the *S. cerevisiae* motif clusters display greater variation in types of clustering. Diversification of motif clusters related to specific cellular processes leads to a more complex structure where multiple inputs are integrated together to regulate target genes for different cellular functions. This is evident in the large motif cluster that contains TUP1 as a central input (Fig. 5). Broadly, this cluster breaks up into three main parts: (top) genes related to mating type switching; (bottom left) genes involved in meiosis; and (bottom right) genes related to biosynthesis and respiration. While each of these cellular processes is controlled in isolation by several independent inputs, more complex phenotypes such as the switching of mating type require the co-ordination of these processes in unison. TUP1 gene fulfills this role and has experimentally been shown to act as a global transcriptional repressor [33]. It has the biological function of switching off genes whose function are not required. The position of TUP1 in the network reflects this role as a repressor of many genes, especially those involved in specialised cell divisions such as meiosis and changes in mating type.

One of the most widely studied examples of an ecological network is the Little Rock Lake food web [22]. Nodes represent species, and edges the consumption of one species by another. Results in Fig. 2c show that most of the FFL clusters in this network (61%) are of types 1 and 2. In these clusters one node is a predator that consumes all or most of the other species in the cluster. The prevalence of these structures reflects the fact that most of the species in the lake are consumed by other species across trophic levels. Only a few top predators exist with most of these acting as omnivores.

The role of omnivory in food webs is still contested [34, 35, 36, 37, 38]. Initial theoretical studies found that its presence under equilibrium conditions can destabilise food webs [34]. However, more detailed collection of species interactions has shown it to be a common property of many types of food web [36]. More recently, non-equilibrium studies of these networks [37, 38] and consideration of potential adaptive mechanisms [37] has revealed that omnivory can help improve system stability and damp potential chaotic dynamics. The FFL motif captures this key relationship and motif clustering the higher-order forms it can take. The prevalence of this and derived structures in the food web is evident with FFL clusters comprising 54.1% of nodes and 38.5% of edges across the entire network (Supplementary Information, Section 2.4). Detailed investigation of other FFL clustering types also showed a decreasing number of longer range predator prey interactions across more distant trophic levels. We found 2- and 3-level predation most common (types 1 and 2), while 4-level relationships were very rare. This is due to the limited number of trophic levels present in this food web. Furthermore, we find it rare for multiple top level predators to share the same low, but alternative intermediate level prey. This is likely due to adaptations to consume one form of prey likely having similar benefits on potential higher-level prey.

In the *C. elegans* neural network 58% of the FFL clusters are also of type 1 or 2. In these networks the nodes are neurons and the edges are synapses. As this is a neural network, type 1 and 2 are clusters where a node receives information from all or most of the other nodes in the cluster. This is consistent with a highly integrated structure with many nodes receiving and integrating information from multiple sources. Furthermore, theoretical studies of the dynamics of FFL structures in neuronal networks have shown their potential role in local stability [39] and permitting input events that do not occur exactly simultaneously to trigger a response through the use of the intermediate node as memory [15]. Apart from these types, others where one and two nodes are shared (2–6 and 7–12) have similar proportions. Kashtan et al. [15] showed that generalised motifs maintain the same function as their underlying motif over a broader range of nodes. Therefore, combining the many functions that motifs have been shown to exhibit, with the ability for neural networks to tune available connection strengths through synaptic plasticity, makes the wide range of motif clustering types in this network potentially capable of many different roles in information processing.

## Engineered and social networks

In contrast, engineered and social networks were dominated by alternative types of motif clustering. The Wikipedia vote and air traffic control networks saw type 6 highly expressed, at 39% and 32% respectively. For the voting network, this reflects the fact that voter preference is likely to match the candidates they vote for. Such relationships arise from the underlying homophily present in virtually all social networks [40, 41, 42] and shown to evolve under many different conditions [43]. While there are unlikely to be direct social ties between voters and candidates, transitivity of this relationship means that candidates voting in additional ballots are likely to match their previous voters choice, embedding type 6 FFL clusters within the network.

Similarly, for the air traffic control network a major influence on the structure is a requirement for robust paths to multiple destinations while taking into account geographical limitations. In this network nodes are airports and edges recommended routes. Types 5 and 6, both highly expressed, embody a function where single input spreads out to many intermediate and output nodes. This suggests that recommended routes attempt to reduce the local burden on specific airports, sharing traffic that may accumulate during difficulties e.g., weather disruptions at particular airports.

The Gnutella network saw a large proportion of types 4 and 12, making up 33% and 18% of the FFL clusters respectively. These types include only a single intermediate node. In this network nodes represent computers (clients or servers) and edges the transmission paths between them. Gnutella is a decentralised file sharing protocol. When a new computer connects to this network and requests a file, it first searches for a local “ultra-peer” server. These are special nodes that act as a high-speed backbone for the network and are purposefully spread out to improve communication efficiency. Once a client is connected, the request is processed and forwarded to the appropriate target server. Once the client is aware of the target it can then directly connect, leading to a FFL transmission structure being generated in the network. Both types 4 and 12 capture this process whereby many clients (inputs) connect to target servers (outputs), using a single ultra-peer server (intermediate).

Finally, the EU e-mail network exhibits larger proportions of type 7 related to multiple inputs and a shared intermediate and output, and type 9 related to a 3-level hierarchy where two intermediate nodes are also connected. The majority of e-mail within an organisation will take place via an organisational hierarchy and these particular motifs seem a likely consequence. One interesting aspect is a lack of other hierarchical motifs being expressed reflecting a segregated structure where it is uncommon for higher level managers to directly interact with those more than two levels below. This would account for type 9 seeing the largest overall expression of 22%, as this type separates the main input from the output through an intermediate layer.

In all cases the real-world networks in Fig. 2c show biases in clustering type distributions that cannot be produced purely by random re-wiring or unbiased node duplication based mechanisms (Fig. 2c; see Supplementary Information, Section 3 for an analytical derivation of the expected motif clustering type distributions for these random models). These distributions suggest that more complex evolutionary mechanisms may be at work such as biased node duplication or strong selective pressures supporting only limited forms of clustering. The motif clustering distributions can be used to uncover biases in these processes (Fig. 3). This is most evident for motif clustering types 7, 10 and 12 (Fig. 2a), which directly relate to duplication of a single input, intermediate, and output node respectively. These are found at elevated proportions when compared to other motif clustering types that share two nodes (types 8, 9 and 11) in many of the biological networks. Furthermore, these duplication based types are not seen in equal proportions. Both the metabolic and transcriptional networks see increased duplication of output nodes which is likely due to the reduced impact this will have on down stream processes during evolution. The Little Rock Lake food web sees an opposite bias towards input duplication/speciation (type 7) which can be accounted for by low-level prey being under increased pressure to diversify so as to evade generalist predators within these structures. In contrast, the neural network does not display any bias towards these three types of clustering, however, node duplication is not known to be a factor in the evolution of this type of network.

**Figure 3.**
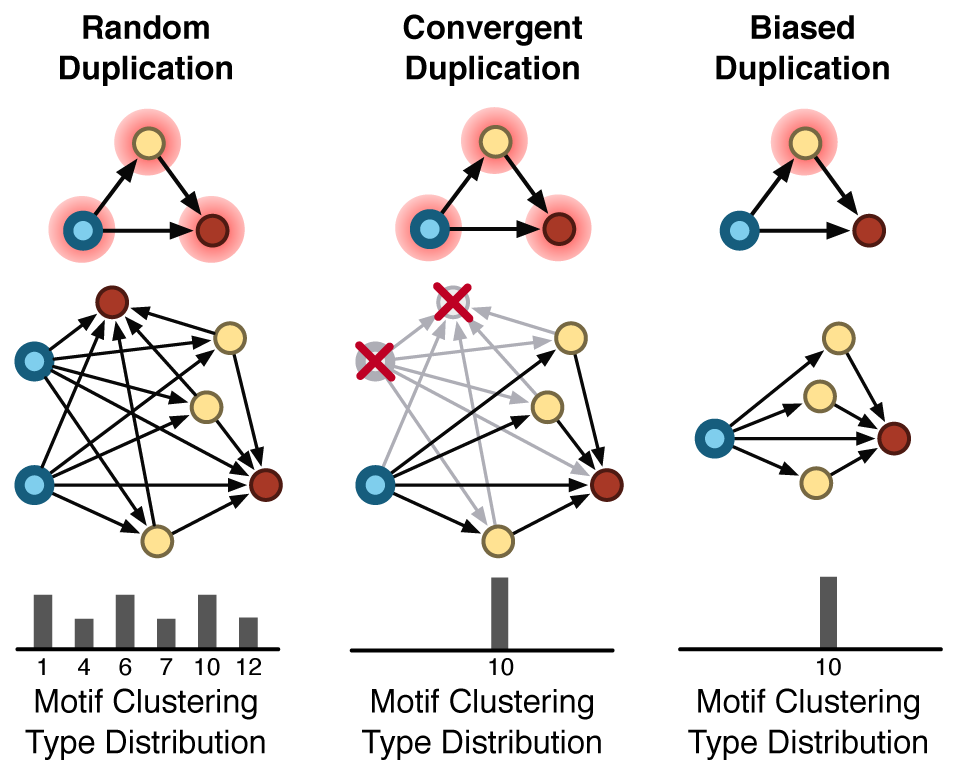
Routes to biased motif clustering distributions for a duplication based growth process. Purely random duplication leads to a broad distribution of specific motif clustering types. In contrast, biased distributions exhibiting a single type of clustering can occur through convergent duplication where all nodes remain equally likely to be duplicated, but post-duplication processes (such as natural selection) remove non-functional or deleterious events. Alternatively, the duplication event itself could be biased, with some types of node having higher probability than others. This bias could be due to functional limitations in the system e.g., physical cabling restrictions in communication networks, or as an intrinsic feature of the system itself e.g., a mutational bias in genetic systems.

To ensure that the motif clustering types were a robust feature, we subjected the networks to random edge removal (Supplementary Information, Section 2.3). In all cases, characteristics of the motif clustering type distributions were maintained for removal of up to 15% of all edges and some networks showed little change even after 50% of edges were removed (e.g., Little Rock Lake food web). Metabolism was found to be the most susceptible to single edge removals, due to the highly interwoven core that is necessary for its carefully controlled function (Supplementary Information, Section 2.3).

## Structure of information flow through the motif clusters

Our results on FFL cluster types suggested that some networks have very strong prevalence for certain types of information flow (according to our results, above 70% of the FFL clusters in a network might be of the same type and hence produce the same type of information flow). Our classification of cluster types identified nodes as input, output or intermediate, but the “intermediate” class encompasses nodes with a range of very different characteristics.

To further dissect information flow within the FFL motif clusters we applied a similar approach to Ma’ayan et al. [10] and used the notion of node spin to classify the extent of each node as a producer, receiver, or relayer of information (Fig. 4a). We compared the node spins of whole networks to those of their FFL clusters, see Fig. 4b (Supplementary Information, Section 2.4). In several cases the FFL subset and the whole network have similar profiles, but there are examples where the FFL subset contributes very differently to information flow from the whole network e.g., metabolism.

**Figure 4.**
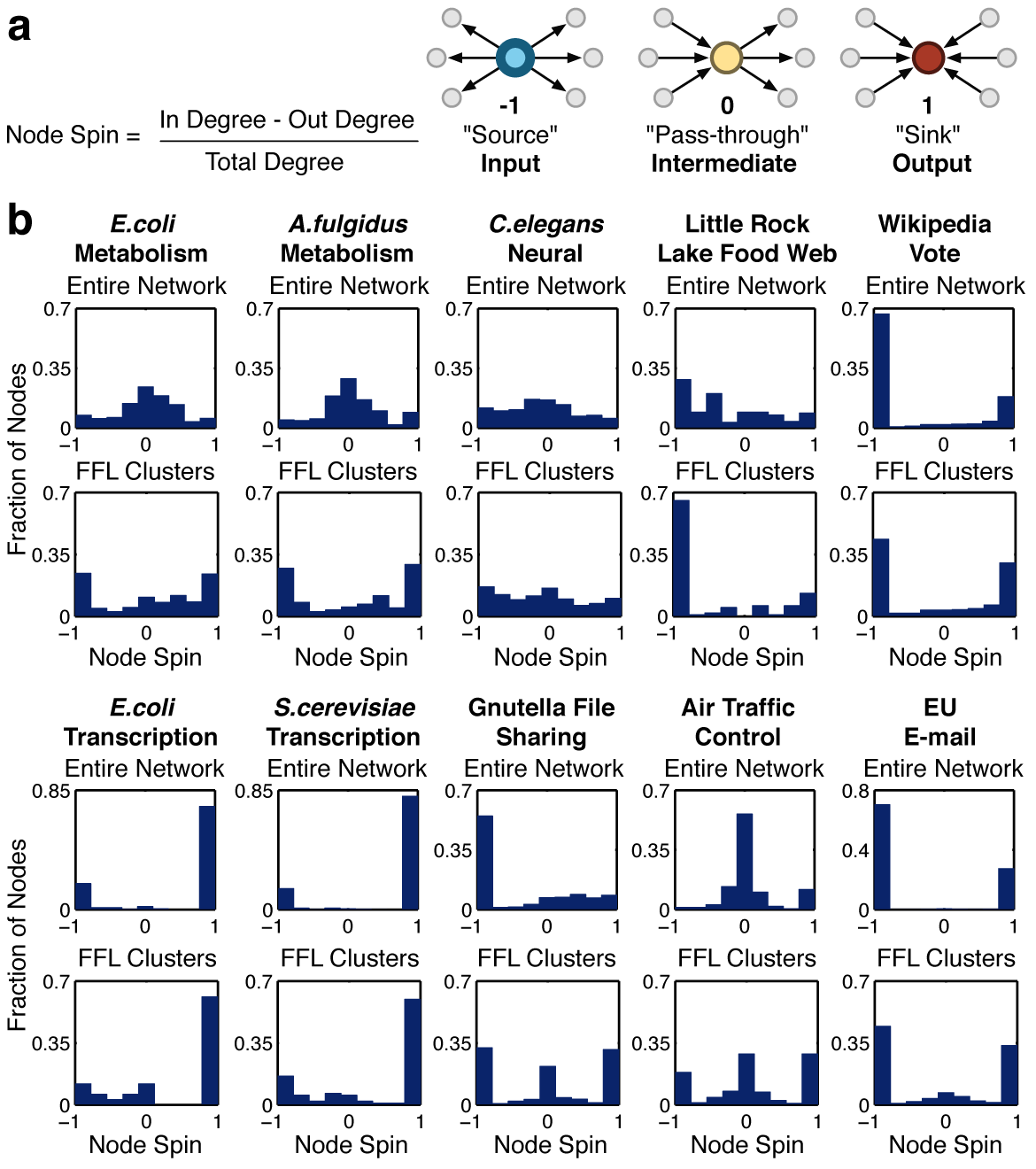
Node spin distributions. (**a**) Node spin is defined as the normalised difference of the in to out degree of a node classifying its role as producing (source), receiving (sink) or relaying (pass-through) information. (**b**) Node spin distributions for the entire real-world networks (top) and the extracted FFL motif clusters (bottom).

Results showed that transcription networks have very polarised node spin distributions. This means that incoming information is integrated through relatively few intermediate nodes before being relayed to large numbers of outputs. This captures the role of master regulators with a few critical signals having to modulate the actions of multiple processes to appropriately alter cellular physiology. In contrast most nodes in metabolic networks have intermediate spins, reflecting their balanced use in multiple biosynthetic pathways.

The incredibly flat distribution of node spins for the C. elegans neural network shows that routes for information flow are evenly distributed with a broad range of input to output ratios. It has been shown that FFLs in these networks can act as decision points, triggering when sufficient numbers of inputs are activated within a short time window [15]. By having a wide-range of potential input-output ratios present in the network, and the previously highlighted ability of these networks to dynamically adapt through synaptic plasticity, they are potentially very flexible to a broad set of alternative functions.

A striking feature of the non-biological networks is the convergent node spin profiles of FFL clusters in the Gnutella file sharing, air traffic control and EU e-mail networks. Even though these are taken from very different domains this convergence reflects the precisely defined roles of components in these human-made networks, where the vast majority of nodes are either solely input, solely output, or solely intermediate. This segregation is generally due to the functional modularity that most man-made systems have. This enables the systems to grow in complexity in a predictable way, with specific tasks encapsulated within particular elements of the system. For example, in the case of the Gnutella file sharing network, inputs and outputs represent the end-users and information providers, while the task of information transmission (intermediate spins) are performed by a specialized set of ultra-peers whose major role is the relay of information. In addition, the transmission of information (Gnutella and EU e-mail) or even planes (air traffic control) relies on the ability of these rely points to not act as potential bottlenecks. This would account for the large proportion of near 0 node spins where a balance of inputs to outputs is present.

Similarly, the Wikipedia vote network displayed a strong convergence to an input/output orientated architecture with a smaller fraction of nodes having an output bias (small ramp in the distributions). This clearly captures the bipartite nature of the voting system, containing members that can be classified as either voters (inputs) or candidates (outputs). Furthermore, as a global feature the extracted FFL clusters display a similar distribution to the entire network and also account for a large proportion of its structure (54.8% of nodes and 85.5% of edges; Supplementary Information, Section 2.4).

## Identifying functionally important nodes in networks

The results discussed above strongly suggest there is a class of nodes that take part in multiple clusters of different types. These nodes are likely to be involved in many parts of a network that carry out different functions. This can make them particularly important for network function and co-ordination. To test this hypothesis we defined a motif clustering type betweenness (MB) measure, see Fig. 5a (Methods). We applied this measure to the S. cerevisiae transcription network (Fig. 5b). The nodes with the highest MB were TUP1, GLN3, GAP1, GAT1 and DAL80. TUP1 is aglobal co-repressor of transcription [44] that switches off well over a hundred genes in diverse signalling pathways. For example, Fig. 5b shows that TUP1 has links to genes involved in mating type switching, meiosis, biosynthesis and respiration. GLN3, GAP1, GAT1 and DAL80 together directly or indirectly regulate hundreds of genes to optimise nutrition [45]. These results confirm that MB can be used to reveal particularly important nodes in networks.

**Figure 5.**
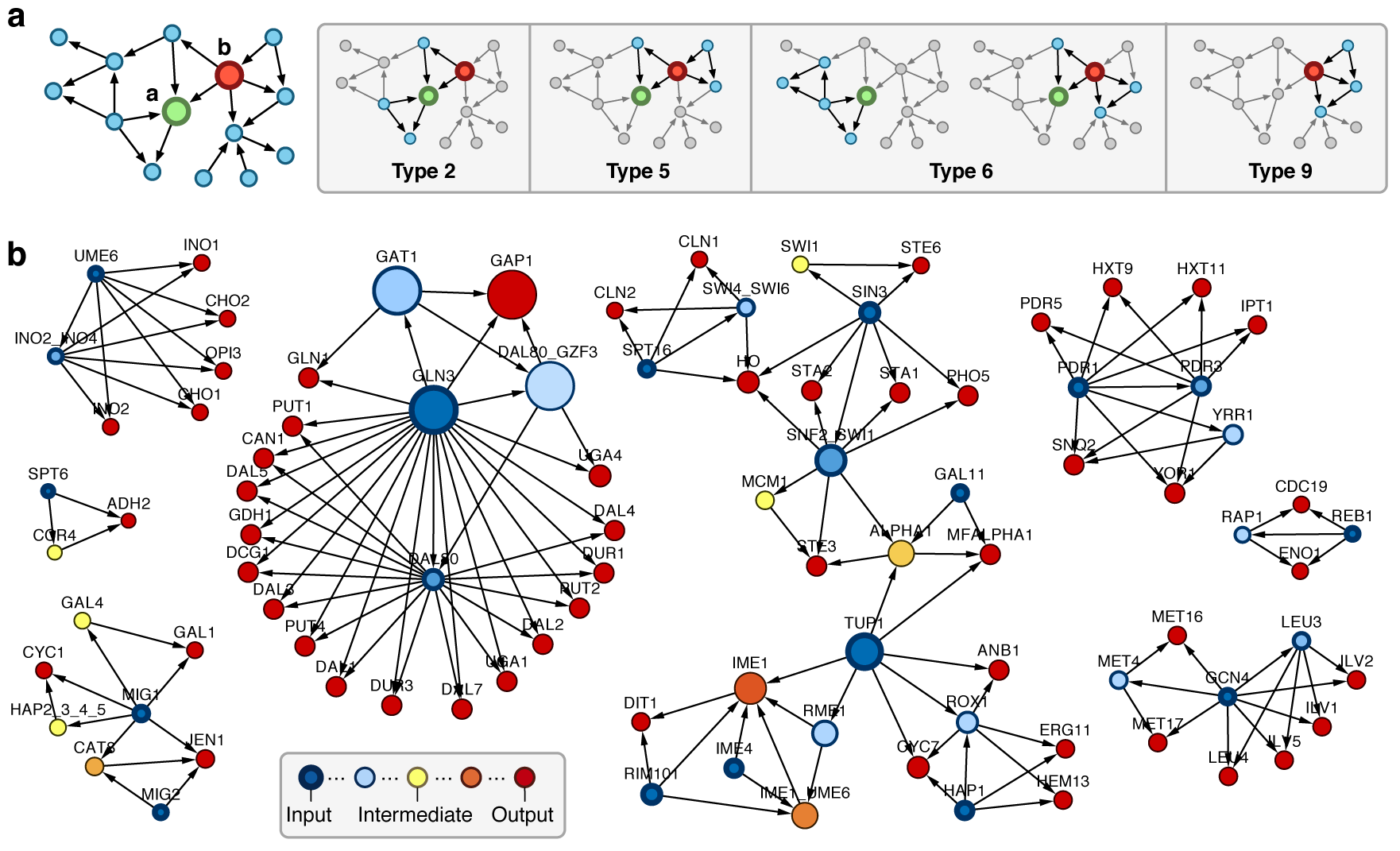
Calculating motif clustering type betweenness (MB). (**a**) Example where two nodes have been selected (left) and highlighted when they are a member of a clustered motif pair (right). The MB is given by the number of different motif clustering types a node is a member of. Therefore, even though both are a member of 4 unique motif pairs, ‘a’ has an MB value of 3 while ‘b’ has a value of 4, due to two motif pairs that include ‘a’ having the same type 6. (**b**) Feed-forward motifs extracted from the S. cerevisiae network. Node size corresponds to MB and colour represents the node spin.

To test this hypothesis further we used experimental data on essential genes in E. coli [46] and assessed whether high MB values of enzymes in the metabolic network were a good predictor of their essentiality (Supplementary Information, Section 4). We compared the accuracy of our predictions using MB to those of other standard measures of node importance, specifically node degree and betweenness centrality (Fig. 6; Supplementary Information, Section 4). While MB displayed excellent performance, a striking difference in its predictions to other measures is shown in Fig. 6 (Supplementary Information, Section 4). The MB predicted nodes exhibiting a broad range of degree and betweenness values which resulted in 3 (∼10%) essential nodes being uniquely identified by MB, see Fig. 6b. This suggests that MB and motif clustering more generally provides a complementary perspective that will help suggest possible essential targets missed by existing approaches.

**Figure 6.**
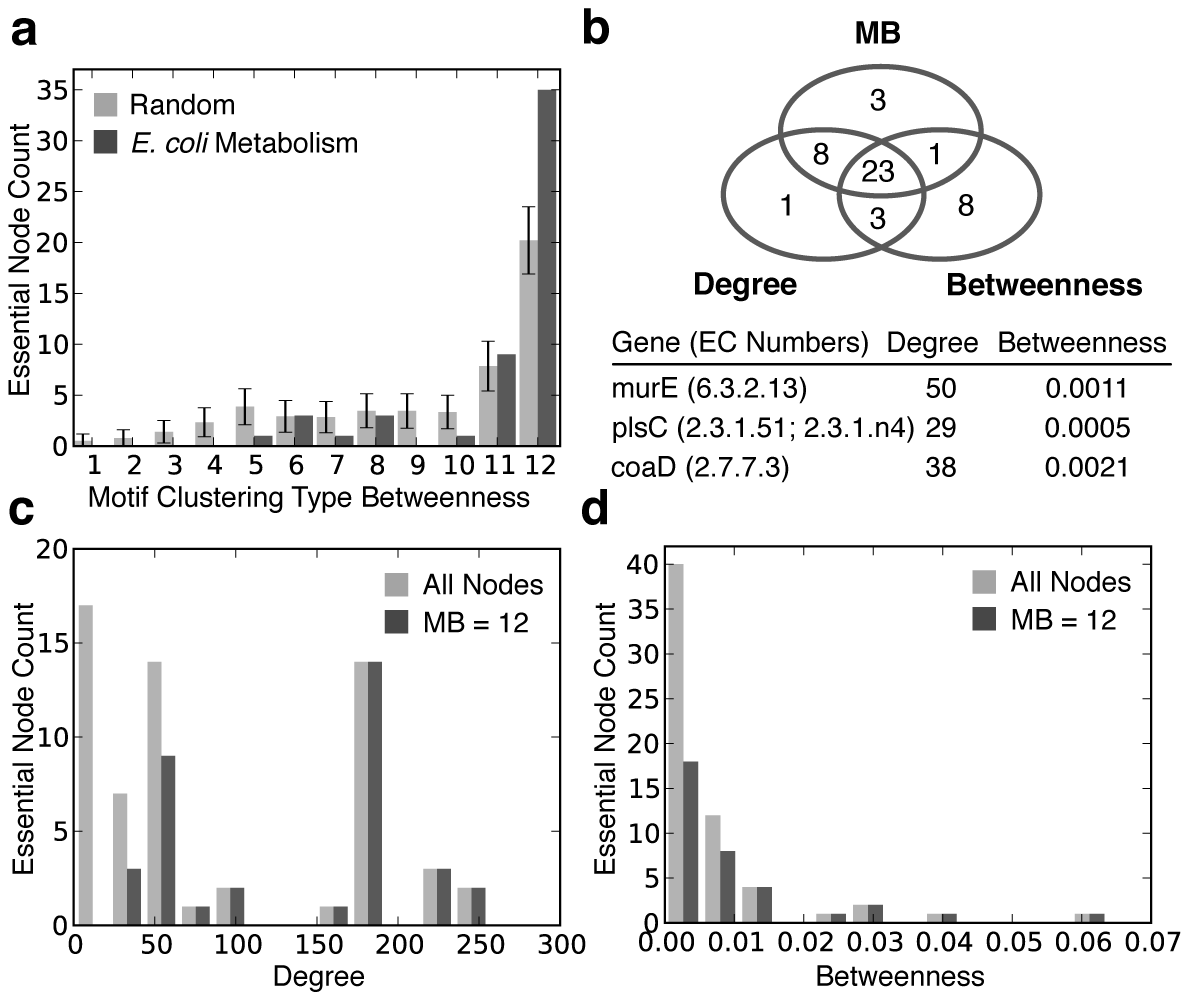
Capturing essential elements of E. coli metabolism with motif clustering. (**a**) Comparison of the number of essential nodes selected for each motif clustering type betweenness (MB) value versus a random selection model that accounts for non-uniform group sizes. (**b**) Venn analysis of essential enzymes selected for the top 156 nodes with greatest importance in regards to our measure i.e., MB = 12. We compared MB, degree and betweenness for all nodes in this set. The three nodes exclusively captured by the MB are displayed in the table below. (**c**, **d**) Comparison of all essential nodes in the metabolic network to all essential nodes with an MB = 12 and the associated (**c**) Degree and (**d**) Betweenness.

It is important to note that while MB works purely using the structure of a network, unlike node based measures such as degree or betweenness, it considers localised structural motifs. The diverse functional dynamics that these motifs in isolation have been shown to exhibit [2, 3, 4, 5, 6, 7] means that analysis of clustering allows for the capture of potential mixing and chaining of differing dynamical functions. Therefore, the MB values are not just a structural measure, but also indirectly incorporate some consideration of potential dynamics or function based on the integration of motifs in different ways. This is a key differentiator of this approach to other topological methods that do not consider these intermediate and functionally relevant structures.

## Discussion

We have explored the way in which motifs can act as building blocks for complex networks and attempted to better understand their connection architecture across several real-world systems. Our analysis revealed that aggregation of FFLs into clusters is significant. Detailed analysis showed that these motifs can potentially become connected in a large number of different ways, but only a few are in fact used within the real-world networks. In some cases just one or two types of cluster dominate. This suggests that limited rules may underly the ways that motifs can be successfully pieced together to generate larger structures with functionally useful dynamics. In addition, biases in the motif clustering types were shown to capture key features of underlying evolutionary processes, such as duplication and divergence, in addition to selective pressures placed upon these processes. Motif clustering offers a glimpse at functionally important structures and the evolutionary process by which they arise. Furthermore, the important role that motif clustering plays was illustrated in terms of the essential nodes within the metabolic network of *E. coli*. The motif clustering type betweenness measure was shown to predict essential nodes with comparable performance to existing approaches while also highlighting nodes missed by other approaches.

The focus of this work has been on motif clustering across entire networks. While this has helped identify system wide architectural principles, many networks are composed of loosely connected communities that often perform distinct functional roles. An intriguing future direction would be to apply these methods at the multiple-resolutions present within a network to capture differences in the motif clustering rules and further refine our understanding of how and why certain types of connection are favoured over others.

## Methods

### Motif Clustering Coefficient

The motif clustering coefficient, *M*_*c*_, attempts to capture the overlap between motifs in terms of the number of shared nodes between all pairs of motif of interest within a network. To calculate the motif clustering coefficient for a network 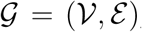, we consider a set of motif types 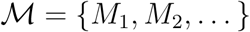, where each 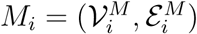 ∀*i*defines a network that captures the motif structure. We also ensure *M*_*i*_ ⊈ *M*_*j*_ ∀*i*, *j* where *i* ≠ *j* such that no motif is a sub-isomorphism of any other. For each motif type 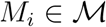 we search for instances (sub-isomorphisms) in 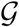 and for each motif occurrence add the associated nodes for the motif from 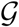 to the set of found motifs *F* = {*f*_1_, *f*_2_,...,*f*_n_}, where 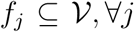. Therefore, *F* contains sets of nodes from 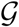 where each set defines one of the motif types in 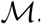 This set was calculated using the VF2 algorithm [47]. The motif clustering coefficient is then given by 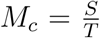 where *S* is the total number of shared nodes between all pairs of found motif in *F*, and *T* is the total possible number of shared nodes had all pairs of these motifs been fully clustered (sharing the maximum possible number of nodes). These are calculated using,

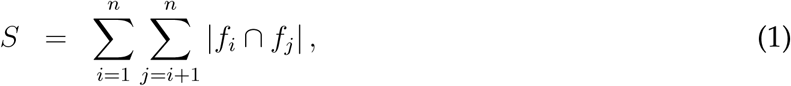

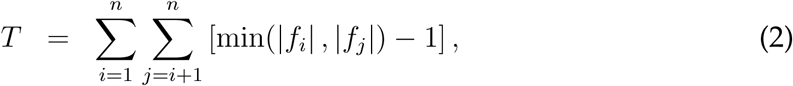

where *n* is the number of motifs in the network, *f*_*i*_ is the set of nodes that make up motif *i*, and | • | denote set cardinality. In the case where only a single type of motif is considered i.e., 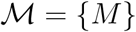 then 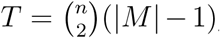, where |*M*| is the number of nodes in motif *M*, and 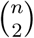 is the binomial coefficient specifying the number of ways of choosing 2 elements from a set of size *n* without taking order into account.

For cases where 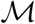 contains several motifs there are two further types of motif clustering we can consider: homologous and heterologous. These are obtained by partitioning the set of found motifs, *F*, as, 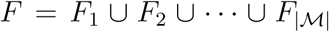, where each subset *F*_*i*_ = {*f*_*i1*_, *f*_*i2*_,…} contains only those nodes that define motifs in the network 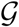 of type *M*_*i*_. Next, we generalise *S* and *T* to consider the number of shared nodes between two specific types of motif,

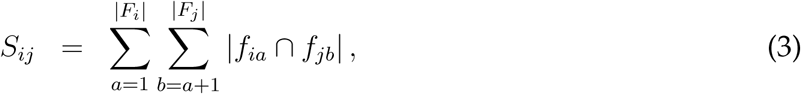

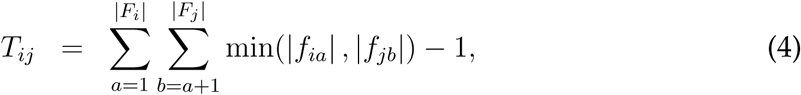

where *S*_*ij*_ is the number of shared nodes between motifs of type *M*_*i*_ and *M*_*j*_, and *T*_*ij*_ is the total possible number shared nodes between the sets of motifs *M*_*i*_ and *M*_*j*_. Then, we are able to consider the clustering between motifs of the same type and define homologous motif clustering as,

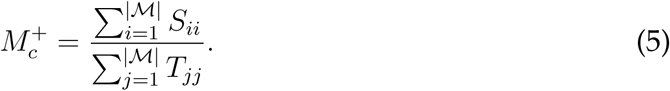

Conversely, by considering the clustering between different types of motif, heterologous motif clustering can be defined as,

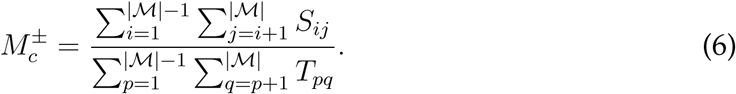

An example for a simple network can be found in Supplementary Information, Section 1.

To calculate the statistical significance of a motif clustering coefficient we used rejection based sampling where random networks were generated maintaining the same number of nodes, edges and motifs as the original. This was performed by starting with an empty network containing the same number of nodes as the original. Motifs were then individually placed at random until the same number of motifs were present. Finally, any outstanding edges were randomly placed to ensure that the same number of edges were also present. If at any point during this process the number of motifs or edges exceeded those found in the original network, the randomised network was rejected and the process restarted. The motif clustering coefficient was calculated for each random network and a comparison made to the original network using a standard Z-score.

### Motif Clustering Type Distributions

To calculate the motif clustering type distribution for a particular set of motif types 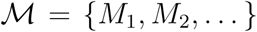, we first generate a set of all possible motif clustering types 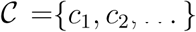 where each member 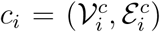, Vi is a network. Specifically, 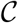 contains networks representing all the unique ways that two motifs from 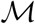 can become clustered sharing at least one node in common. To do this we enumerate over all possible overlaps between all motifs in 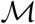 including the case where a particular motif is clustered with itself e.g., if 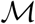 contained a single feed-forward loop motif then the set of motif clustering types *C* would contain all the networks shown in Fig. 2a. When generating this set, it is important to ensure all clustering types are unique by checking that any newly generated candidates are not isomorphic to an existing member of *C*. This ensures symmetries in a motif do not lead to multiple clustering types that have the same network structure. Once the set *C* of motif clustering types has been generated, we search for all instances of the motif types in *M* within the network of interest 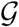 using the same approach as for the motif clustering coefficient. For all pairs of motif found, we extract them from the original network and compare the resultant sub-graph to the set of motif clustering types in *C*. Cases where motifs do not share any nodes are neglected. We count the occurrences of each motif clustering type and normalise by the sum total of all motif clustering type counts. As with the motif clustering coefficient, this measure is easily adapted to homologous and heterologous cases by generating motif clustering types C that only enumerate clustering between the same or different types of motif in 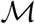 respectively.

### Motif Clustering Type Betweenness

The motif clustering type betweenness of a node is calculated by extracting all motifs of interest that contain the selected node as a member. We then consider all pairs of these motifs and for each classify their type in *C* using the same method as described for the motif clustering type distributions. The motif clustering type betweenness measure is then given by the number of different types of motif clustering in *C* that the node is a member of. An example can be found in Fig. 5a.

## Acknowledgements

We would like to thank Peter Green and Ayalvadi Ganesh of the University of Bristol, UK and Mauricio Barahona of Imperial College London, UK for their insightful discussions on this work. T.E.G., C.S.G. and M.B. were supported by BrisSynBio, a BBSRC/EPSRC Synthetic Biology Research Centre (grant BB/L01386X/1). T.E.G. also acknowledges support of EPSRC-GB Grant No. EP/E501214/1. Analysis was carried out using the computational facilities of the Advanced Computing Research Centre, University of Bristol, UK – http://www.bris.ac.uk/acrc.

### Author Contributions

All authors conceived and designed the study, analysed the data, and wrote the manuscript. T.E.G. performed the numerical experiments.

### Competing financial interests

The authors declare no competing financial interests.

